# Host traits and environmental variation shape gut microbiota diversity in wild threespine stickleback

**DOI:** 10.1101/2024.11.19.624388

**Authors:** Andreas Härer, Emma Kurstjens, Diana J. Rennison

## Abstract

**Background:** Despite the growing recognition of the importance of gut microbiota in host ecology and evolution, our understanding of the relative contributions of host-associated and environmental factors shaping gut microbiota composition within and across wild populations remains limited. Here, we investigate how host morphology, sex, genetic divergence, and environmental characteristics influence the gut microbiota of threespine stickleback fish populations from 20 lakes on Vancouver Island, Canada.

**Results:** Our findings reveal substantial variation in gut microbiota composition and diversity among populations, with host traits exerting a stronger influence on bacterial alpha diversity than environmental characteristics. Host morphology, which is indicative of trophic ecology, was linked to gut microbiota divergence among populations, suggesting that dietary specialization may play a role in shaping the stickleback gut microbiota. Within and across populations, we only observed a weakly defined core microbiota and limited sharing of ASVs among hosts, indicating that gut microbiota composition is individualized. Additionally, we detected sex-dependent differences in microbial diversity, opening avenues for future research into the mechanisms driving this variation.

**Conclusions:** In sum, our study emphasizes the need to consider both host-associated and environmental factors in shaping gut microbiota dynamics and highlights the complex interplay between host organisms, their associated microbial communities, and the environment in natural settings. Ultimately, these insights enhance our understanding of host-microbiota interactions and their eco-evolutionary implications.

## Background

The gut microbiota plays a vital role in many aspects of an organism’s biology, which can have eco-evolutionary implications (Suzuki 2017, Gould et al. 2018, Rudman et al. 2019, Moeller and Sanders 2020, Henry et al. 2021). Understanding the dynamics that shape gut microbiota of animals is therefore of particular importance in wild populations, where organisms are exposed to complex environmental pressures and interactions that are often absent in controlled settings. Numerous studies across a broad range of study systems have highlighted the multifaceted influences of both host traits and environmental factors on gut microbiota composition and diversity. Key host traits include genetics (Benson et al. 2010, Bulteel et al. 2021), immune system function (Hooper et al. 2012, Bolnick et al. 2014a), sex (Martin et al. 2010, Stoffel et al. 2020), and trophic ecology (Delsuc et al. 2014, Baniel et al. 2021). Environmental factors include habitat type (Shankregowda et al. 2023), physicochemical characteristics (Bestion et al. 2017, Fontaine et al. 2018), geographic context (Moeller et al. 2013, Moeller et al. 2017), and interaction with other hosts (Li et al. 2016, Burns et al. 2017). Studying multiple key factors within a single system and thereby disentangling their relative contributions can be challenging, but it offers the potential to reveal new insights into how gut microbial communities are structured across wild populations. Host species that have recently and repeatedly colonized distinct habitats and adapted to local conditions can offer valuable insights into the factors driving gut microbiota variation across ecologically and genetically divergent, geographically isolated natural replicates.

The threespine stickleback fish (*Gasterosteus aculeatus*, hereafter referred to as ‘stickleback’), a well-known model system for ecology and evolution (Bell and Foster 1994), represents such a system. Since the end of the last ice age (10,000-12,000 years ago), marine stickleback independently colonized and adapted to thousands of freshwater habitats. Among these freshwater populations there is substantial ecological, morphological, and genetic divergence (Schluter and McPhail 1992, Schluter 1993, Bell and Foster 1994, Matthews et al. 2010, Rennison et al. 2020). A particularly important axis of divergence is reflected in their trophic ecology. Freshwater stickleback typically consume two distinct types of prey found in different habitats; benthic prey, which are littoral invertebrates found in the sediment, and limnetic prey, which are pelagic zooplankton found in the open water of lakes (Bell and Foster 1994). Variation in the proportion of these diet types has been detected within and across lakes, and stickleback’s trophic ecology ranges from benthic specialists to generalists to limnetic specialists (Schluter and McPhail 1992, Bolnick and Ballare 2020). Variation in trophic ecology is associated with stickleback morphology in terms of overall body shape as well as the feeding apparatus and body armor (Schluter 1993, Bell and Foster 1994, McGee et al. 2013). Given this remarkable ecological and morphological diversity, stickleback are instrumental for investigating gut microbiota dynamics and the interplay between the environment, host organisms and their associated microbial communities.

Stickleback are also emerging as a model system for gut microbiota research, and initial work has greatly advanced our understanding of how host traits and environmental factors contribute to gut microbiota variation (Bolnick et al. 2014b, Bolnick et al. 2014c, Milligan-Myhre et al. 2016, Rennison et al. 2019, Steury et al. 2019, Härer and Rennison 2023, Small et al. 2023, Härer and Rennison 2024). Previous work has indicated that host traits affecting gut microbiota composition include sex (Bolnick et al. 2014c), diet and ecotype (Bolnick et al. 2014b, Smith et al. 2015, Rennison et al. 2019, Härer and Rennison 2024), and genomic variation (Smith et al. 2015, Steury et al. 2019, Small et al. 2023). These studies have laid an essential foundation for our understanding of the stickleback gut microbiota. However, they come with certain limitations for advancing our understanding of the factors that shape ecologically relevant gut microbiota variation within and across wild host populations. For example, previous studies were either restricted to a single or few lake populations (Bolnick et al. 2014a, Bolnick et al. 2014b, Bolnick et al. 2014c, Smith et al. 2015, Milligan-Myhre et al. 2016, Steury et al. 2019), included very few individuals per populations hindering the assessment of within-population diversity (Rennison et al. 2019), or were conducted in the lab (Härer and Rennison 2023, Small et al. 2023).

Building on the progress made in earlier studies, we sought to explore host-microbiota interactions more comprehensively in wild populations. To this end, we surveyed populations from twenty lakes on Vancouver Island, Canada (Figure 1), to investigate how host sex, morphology (a proxy for trophic ecology), and genetic differentiation, in addition to each lake environment’s physicochemical characteristics and bacterial communities, and geographic distance from each other, contribute to shaping the stickleback gut microbiota. Specifically, we tested the effects of these factors on gut microbiota alpha diversity (bacterial diversity of individual hosts) and beta diversity (dissimilarity of bacterial communities among hosts) across populations, as well as the effects of host morphology and sex on alpha and beta diversity within populations. In this study, we aimed to synthesize these diverse influences within a single framework to unravel their relative contributions to gut microbiota composition and diversity, which can advance our understanding of the complex dynamics governing microbial communities within the gut. By integrating these factors and exploring their interplay, we seek to uncover novel insights into the structuring of the gut microbiota and the geographic distribution of microbial diversity across wild populations of freshwater fish.

**Figure 1:**
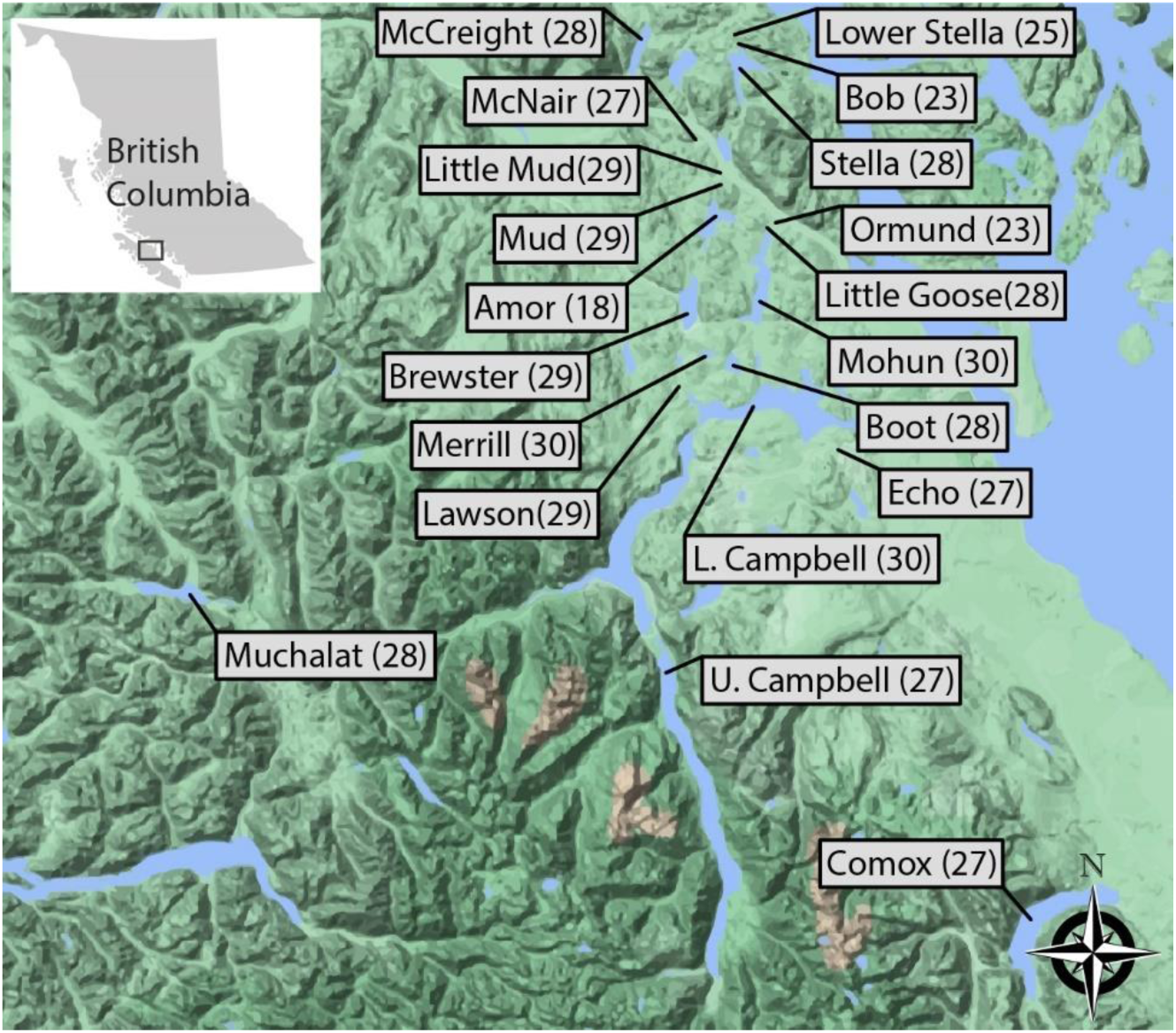
Map of Vancouver Island, Canada, showing locations of the 20 lakes included in this study. For each lake the sample size for the corresponding stickleback population is indicated. The lakes are distributed across several watersheds and this map provides an overview of the geographic distribution of the sampled populations. This data allows investigating how ecological, geographic, and environmental factors influence the stickleback gut microbiota.

## Methods

### Sampling and data collection

We collected stickleback using minnow traps in the spring of 2020, 2021, and 2022 under British Columbia Fish Collection permits NA20-602264, MRVI21-619908, and NA22-713085, respectively. In the field, we euthanized fish with an overdose of MS-222 (500 mg/L) and stored them at −20°C until gut dissection. In the same year of fish sampling, we further collected four water samples from the shoreline of each lake. Precautionary measures were taken to mitigate contamination risk across lakes. We sterilized sampling equipment (forceps, syringes, filter housings, and 1.5 mL tubes) using UV light before the field trip and we assembled individual sampling kits for each lake. These kits were kept sealed until sampling at the designated location. We used disposable gloves and avoided direct contact with the water to prevent human contamination. Each sample measured 120 mL and was manually filtered through a cellulose nitrate filter (Whatman plc, Maidstone, UK; ø 25 mm, pore size 0.2 µm) using a 50 mL sterile syringe equipped with a Luer lock. Following filtration, we carefully removed the filters from the housing using sterile disposable forceps and transferred them into 1.5 mL tubes. These tubes were then stored at −20°C until DNA extraction.

To determine body shape of all fish included in our study, we used geometric morphometric measurements of stickleback body shape based on 17 digital landmarks (Albert et al. 2008). To collect this data, we took standardized photographs of each specimen’s left lateral side including a ruler for scale. We used tpsDig2 (Rohlf 2006) for landmarking and imported data into morphoJ (Klingenberg 2011) for Procrustes fitting, principal component analysis (PCA), and creating a covariance matrix. We then imported these files into R v4.2.1 (R Core Team 2022) where we created Mahalanobis distance matrices on the individual and population level (*mahalanobis* function of the stats package v4.2.1) that were used for subsequent analyses. Additionally, we obtained pairwise F_ST_ values from Bolnick and Ballare (2020) for sixteen of the populations included in our study to determine levels of genetic differentiation among populations. Please note that these F_ST_ values are based on stickleback collected in the same lakes but in different years compared to our samples. Geographic distances between lakes were measured as straight-line paths using the closest points along their shorelines, as determined with the distance measurement feature in Google Maps (Google Maps). In the spring of 2022, we collected various physicochemical characteristics of the lakes (temperature, dissolved oxygen, conductivity, pH) using a YSI Professional Plus device (YSI Incorporated, Yellow Springs, OH). For each lake, we collected three replicate measurements and averaged the values.

### Sample processing and library preparation

In the lab, we rinsed fish with EtOH and dissected their whole guts using sterile equipment. To minimize the abundance of transient bacteria, we carefully removed gut contents by squeezing. We then stored gut samples at −80°C until DNA extraction. We extracted DNA from fish guts with the QIAGEN PowerSoil Pro Kit according to the manufacturer’s protocol (Qiagen, Hilden, Germany) and from cellulose nitrate filter with the QIAGEN DNeasy Blood & Tissue kit (Qiagen, Hilden, Germany) with minor modifications. As a first step, we immersed filters in 450 µl buffer ATL and 50 µl proteinase K and incubated them at 65°C for 1h. Then, we added 500 µl buffer AL and 500 µl 100% ethanol and rigorously vortexed the tubes. Next, we applied the mixture to a DNeasy Mini spin and performed the rest of the extraction according to the manufacturer’s protocol. DNA extractions were done under sterile conditions in a laminar flow hood.

To characterize the taxonomic composition of host-associated and free-living bacterial communities, we used a 16S rRNA gene metabarcoding approach. Specifically, we amplified the V4 region of the 16S rRNA gene with barcoded 515F and 806R primers (for information on primer and barcode sequences see https://github.com/SchlossLab/MiSeq_WetLab_SOP/blob/master/MiSeq_WetLab_SOP.md). We conducted PCR amplification in triplicate, using a 10 μl reaction volume and the Platinum II Hot Start PCR Master Mix (Thermo Fisher Scientific). Subsequently, we pooled the three replicates. We included negative controls of sterile water during DNA extraction and PCR amplification, which did not yield measurable DNA concentrations. The PCR protocol involved an initial denaturation step for 60 s at 98 °C, followed by 35 amplification cycles with 10 s at 98 °C, 20 s at 56 °C, and 60 s at 72 °C, and a final elongation step for 10 min at 72 °C. We visually confirmed amplification of fragments of expected size by gel electrophoresis (2% agarose gels) and we measured DNA concentrations of amplicons with a Qubit 4 Fluorometer (Thermo Fisher Scientific, Waltham, MA). Finally, we pooled samples in an equimolar manner, and the final libraries were sequenced on the Illumina MiSeq 600 (PE300) platform at the UC Davis Genome Center after bead clean-up and quality check on a Bioanalzyer (Agilent Technologies, Santa Clara, CA). Unfortunately, DNA extraction and amplification failed for water samples from Ormund Lake, leaving us with nineteen lakes for which we have information for host-associated and free-living bacterial communities.

### Data analysis

The sequencing runs produced a total of 21,912,056 raw sequencing reads, with mean values of 31,243 reads per stickleback gut and 56,719 reads per water sample (Table S1). We imported sequence data into QIIME2 (Bolyen et al. 2019) and due to low sequencing depths for some samples we used 250 bp of the forward reads (which constitutes 86% of the amplified region) for our analyses because merging of reads further reduced read numbers. Briefly, we used the QIIME2 plugin *dada2* for sequence quality check, read correction, and chimera filtering in order to obtain amplicon sequencing variants (ASVs) (Callahan et al. 2016). We then constructed a bacterial phylogeny with FastTree 2.1.3 (Price et al. 2010) and assigned taxonomy according to the SILVA 138 ribosomal RNA (rRNA) database at a 99% similarity threshold (Quast et al. 2013). Finally, we removed chloroplasts, mitochondria, and archaea and further filtered ASVs that were found only in a single sample and had less than 10 reads or could not be assigned below the phylum level. After filtering, we retained an average of 18,611 reads per stickleback gut and 24,422 reads per water sample (Table S1). We normalized our ASV table through scaling with ranked subsampling (SRS) with a C_min_ of 2500 reads (Beule and Karlovsky 2020), which left us with a total of 617 and sample sizes per lake ranged from 23-30 for stickleback guts (except for Amor Lake for which sample size was only 18) and from 3-4 for water samples (Table S2). We visualized taxonomic composition of the gut microbiota for each stickleback population on the levels of bacterial phylum level; only phyla comprising more than 2% of the microbiota in any of the populations are shown (Figure 2A).

**Figure 2:**
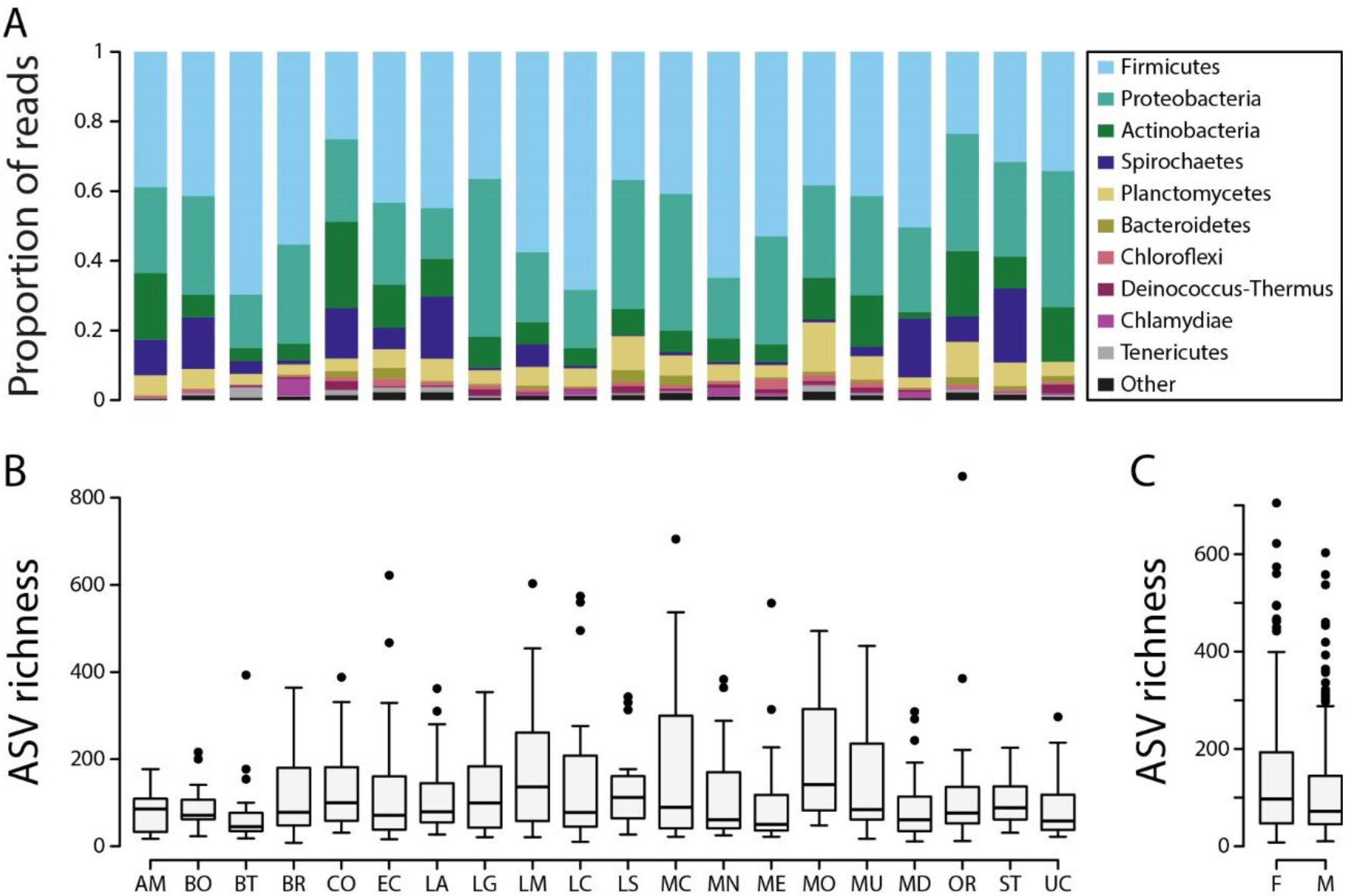
Taxonomic bar plots on the phylum level summarized by population (A). Alpha diversity, shown as ASV richness, by population (B) and by host sex (C).

We used three metrics to describe alpha diversity (bacterial diversity of individual hosts): ASV richness, Shannon diversity, and Faith’s phylogenetic diversity, and three metrics to describe beta diversity (dissimilarity of bacterial communities among hosts): Bray-Curtis dissimilarity, unweighted and weighted UniFrac (Lozupone and Knight 2005, Lozupone et al. 2011). Across all populations, we used negative binomial generalized linear mixed-effects models (g*lmer.nb* function in the ‘lme4’ package v1.1-31) (Venables and Ripley 2002) for ASV richness to account for skewness and overdispersion of count data. For Shannon diversity and Faith’s phylogenetic diversity we used linear mixed-effects models (*lmer* function in the ‘lme4’ package v1.1-31). All models include lake as random effect and host sex, the first three principal components obtained from geometric morphometric measurements of stickleback body shape, as well as temperature, dissolved oxygen, conductivity, and pH of the lake environment as fixed effects. We then used type III ANOVA to test for statistical significance of the model terms (*Anova* function in the ‘car’ package v3.1-0). To contrast alpha diversity between sexes, we used two sample t-tests (*t.test* function in the ‘stats’ package v4.2.1). We further tested for effects of host sex and the first three geometric morphometrics principal components on the three alpha diversity metrics separately for each population, P values were adjusted for multiple testing using FDR correction.

To test for dissimilarity of microbial communities, we used PERMANOVA (*adonis2* function in the ‘vegan’ package v2.6-2) (Anderson 2001, Oksanen et al. 2019). First, we ran PERMANOVA with lake and sample type as independent variables to determine whether bacterial community composition is mainly structured geographically across lakes or between host-associated and free-living communities (fish guts vs lake water). Next, we ran PERMANOVA with host sex, the first three geometric morphometrics principal components, as well as temperature, dissolved oxygen, conductivity, and pH of the lakes as independent variables. Similar to the alpha diversity analyses, we also tested for the effects of host sex and the first three geometric morphometrics principal components on the three beta diversity metrics separately for each population and P values were adjusted for multiple testing using FDR correction. To study the core gut microbiota, we identified the ASVs shared among 50% and 80% of hosts and assigned their taxonomy on the bacterial family level. We conducted these analyses within each population and also across all populations. Additionally, we calculated within-population beta diversity for each lake to obtain information on how dissimilar hosts of the same population are. Moreover, we generated population-level distances matrices and used multiple regression on distance matrices (MRM), an extension of partial Mantel analysis, to test for effects of divergence in free-living bacterial communities of the lakes, host morphology and genetics, as well as geographic distance on the three gut microbiota beta diversity metrics (*MRM* function in the ‘ecodist’ package v2.1.3) (Goslee and Urban 2007). For this analysis, we only used a subset of 15 lakes for which we had all the data mentioned above.

## Results

The taxonomic composition of stickleback gut microbiota revealed both commonalities and variation across populations. While the major bacterial phyla were shared, their relative abundances varied among populations (Figure 2A). The most abundant phylum was Firmicutes, accounting for an average of 44.8% of sequencing reads, with population averages ranging from 23.5% to 69.6%. This was followed by Proteobacteria at 27.2% (ranging from 14.6% to 45.4%), Actinobacteria at 10% (ranging from 2% to 24.8%), Spirochaetes at 6.4% (ranging from 0.1% to 21.3%), and Planctomycetes at 5.8% (ranging from 2.9% to 14.1%) (Figure 2A). These differences in relative abundances underscore the dynamic nature of the stickleback gut microbiota across wild populations, likely reflecting both ecological and evolutionary processes. This variation offers a unique opportunity to investigate the factors that explain the observed differences in bacterial community composition.

### Across- and within-population variation in alpha diversity

Alpha diversity of the stickleback gut microbiota exhibited considerable variation across populations. Mean alpha diversity on the population level ranged from 69.5 to 205.7 for ASV richness (Figure 2B), from 2.5 to 4.1 for Shannon diversity and from 9.2 to 19.4 for Faith’s phylogenetic diversity. All three alpha diversity metrics were associated with host sex (ASV richness: *χ^2^* = 11.18, *P* < 0.001; Shannon diversity: *χ^2^* = 2.74, *P* = 0.098; Faith’s phylogenetic diversity: *χ^2^* = 10.08, *P* = 0.002) and the second principal component of geometric morphometric analysis (ASV richness: *χ^2^* = 6.17, *P* = 0.013; Shannon diversity: *χ^2^* = 3.89, *P* = 0.048; Faith’s phylogenetic diversity: *χ^2^* = 4.84, *P* = 0.028). Pairwise tests revealed that females had 24.4% higher ASV richness (*t* = 2.72, *P* = 0.007; Figure 2C), 6.2% higher Shannon diversity (*t* = 1.82, *P* = 0.069) and 16.4% higher Faith’s phylogenetic diversity (*t* = 2.67, *P* = 0.008) compared to males. Yet, no significant effects of environmental parameters on any of the alpha diversity metrics were detected.

When analyzing populations separately, we found little evidence for effects of host sex and morphology on alpha diversity after FDR correction. ASV richness was only associated with host sex in Lower Campbell Lake (*χ^2^* = 11.57, *P* = 0.013) and potentially Comox Lake (*χ^2^* = 6.87, *P* = 0.088). The first geometric morphometrics principal component was associated with ASV richness in Upper Campbell Lake (*χ^2^* = 11.16, *P* = 0.017), and the second geometric morphometrics principal component was associated with ASV richness in Lower Campbell Lake (*χ^2^* = 10.35, *P* = 0.013) and McNair Lake (*χ^2^* = 12.23, *P* = 0.009) (Table S3). For Shannon diversity, there was only suggestive evidence for an effect of host sex (*F* = 10.41, *P* = 0.078) and the first geometric morphometrics principal component (*F* = 9.77, *P* = 0.098) in Comox Lake (Table S4). No within-population effects of host sex and morphology were found for Faith’s phylogenetic diversity (Table S5).

### Within- and across-population core microbiota and within-population gut microbiota dissimilarity

Across all populations, we found little evidence for a core gut microbiota. On average each ASV was shared among 3.2 hosts, which represents 0.6% of all hosts surveyed in this study. Further, a large proportion of ASVs, 13,925 out of 21,263 representing 65.5%, were found only in a single host. Only four ASVs were shared among more than 50% of all hosts; two of these ASVs belonged to the bacterial family Halomonadaceae, and the other two belonged to the Burkholderiaceae and Mycobacteriaceae. No ASV was shared among more than 80% of all hosts. Next, we determined the core gut microbiota on the population level with mixed results across populations: the number of ASVs shared among 50% of hosts ranged from four in Lawson Lake to sixteen in McNair Lake whereas the number of ASVs shared among 80% of hosts ranged from zero in four lakes (Lawson, McCreight, Ormund, and Stella) to three in five lakes (Comox, Little Mud, McNair, Merrill, and Muchalat) (Figure 3A). At the 50% threshold we found a total of 50 core ASVs; 22 of them were part of the core microbiota only in one of the populations. Yet, six ASVS were part of the core microbiota in more than ten populations and two of these each belonged to the bacterial families Burkholderiaceae and Halomonadaceae and one each belonged to the Bacillaceae and Mycobacteriaceae. Notably, one of the Halomonadaceae ASVs was part of the core microbiota in all twenty populations. At the 80% threshold, we only found eleven core ASVs; nine of which were part of the core in one or two populations. Only two ASVs were part of the core microbiota in seven populations and these ASVS belonged to the families Burkholderiaceae and Halomonadaceae. Moreover, there was little variation in the number of hosts each ASV was shared among with average values ranging from 1.5 to 1.8 individuals, representing 5% to 8.5% of the populations (Figure 3A). This weak evidence for a core microbiota on the population level was corroborated by relatively high levels of within-population beta diversity across all lakes (Figure 3B). Based on the three beta diversity metrics, values ranged from 0.92-0.95 (Bray-Curtis dissimilarity), 0.73-0.76 (unweighted UniFrac), and 0.55-0.67 (weighted UniFrac). In sum, most ASVs were found only in one or very few hosts and only a small number of ASVs was found in the majority of hosts within each population.

**Figure 3:**
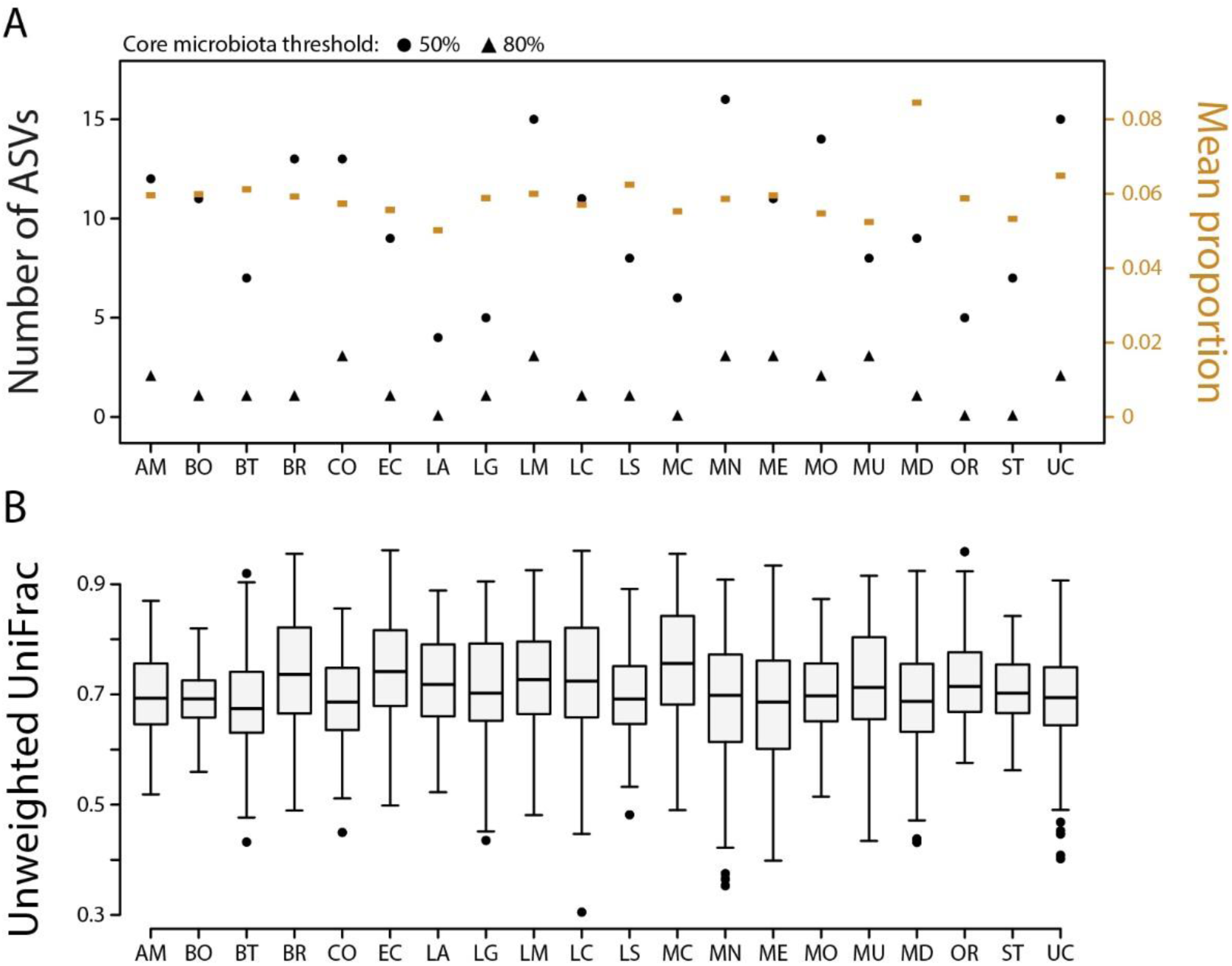
Core gut microbiota (A) and within-population beta diversity across stickleback populations (B). The number of core ASVs at the 50% threshold ranged from 4 to 16 across lakes, while at the 80% threshold, it ranged from 0 to 3 (A). On average, ASVs were shared among 5% to 8.5% of individuals within a population (A). Within-population beta diversity was consistently high across lakes based on unweighted UniFrac, with average values ranging from 0.73 to 0.76 (B). Together, these results demonstrate limited evidence for a core microbiota and high within-population variability across wild stickleback populations.

### Gut microbiota dissimilarity on the individual host level (within- and across-population beta diversity)

We found that microbial communities from stickleback guts and lake water differed substantially (Figure 4). Next, we focused specifically on the gut microbiota and determined the main contributors to bacterial community dissimilarity across stickleback populations by testing host-associated (sex, morphology) and environmental (temperature, dissolved oxygen, pH) factors. While all the tested factors affected gut microbiota dissimilarity, none of them explained more than 1% of the variation in beta diversity (except for temperature which explained 1.1% of the variation based on weighted UniFrac). All these factors combined explained 3.7-5.8% of the variation in beta diversity based on the three metrics (Table S6).

**Figure 4:**
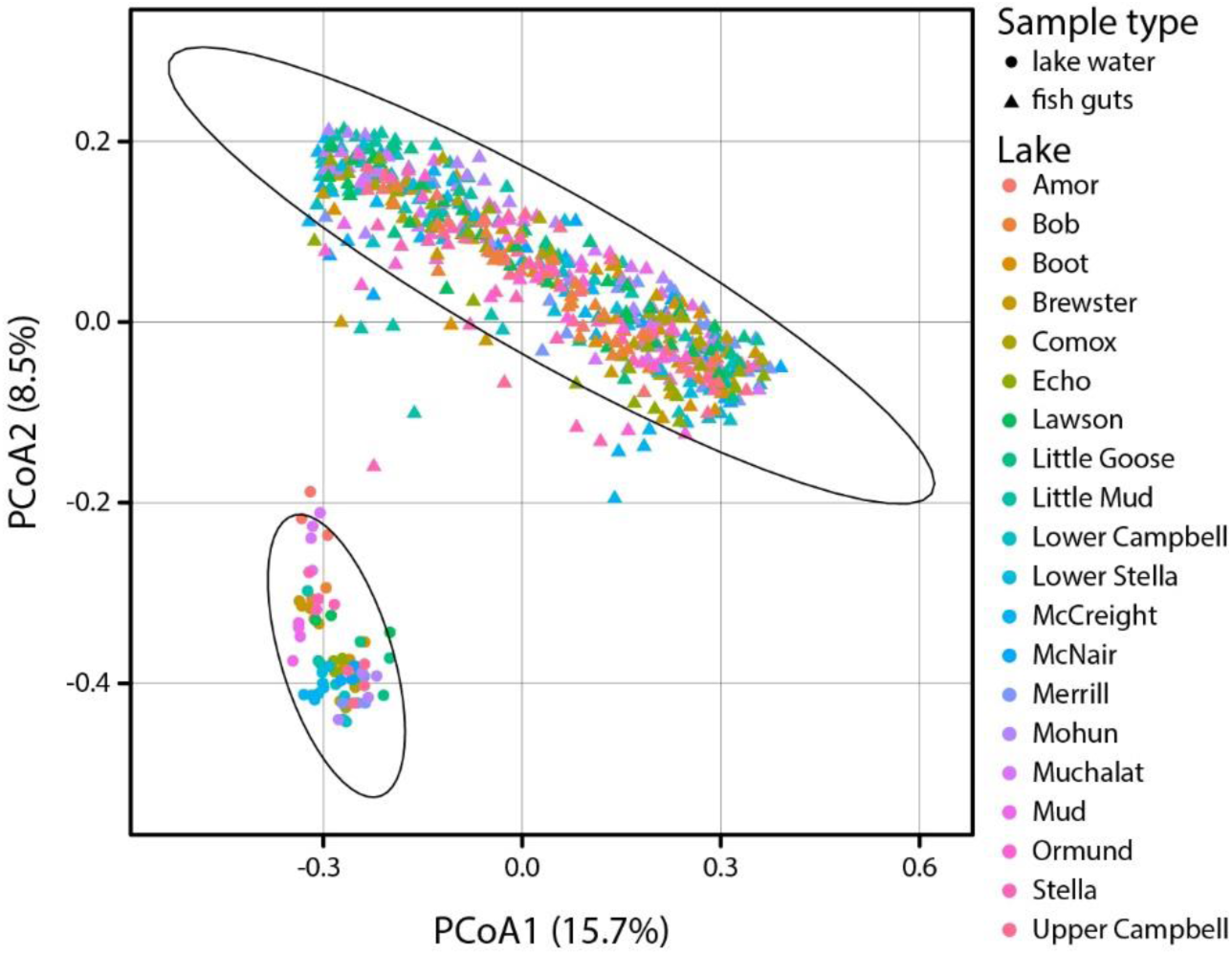
Principal Coordinates Analysis (PCoA) plot based on the unweighted UniFrac metric illustrating the variation in microbial community composition among fish guts (triangles) and water samples (circles) collected from different lakes. Each point represents a sample with colors indicating the lake of origin. The axes correspond to the first two principal coordinates, which capture the majority of the variance in the data. Ellipses denote the 99% confidence intervals for the clustering of sample types (fish guts and lake water).

When testing beta diversity among individuals of each population separately, we obtained similar results as for alpha diversity: only in a few populations did we find evidence for associations between host sex or morphology and beta diversity after FDR correction. We only found suggestive detected for effects of the first geometric morphometrics principal component in Merrill Lake (*R^2^* = 0.079, *F* = 2.487, *P* = 0.06) and host sex in Bob Lake (*R^2^* = 0.1, *F* = 2.347, *P* = 0.08) and Mohun Lake (*R^2^* = 0.056, *F* = 1.654, *P* = 0.08) based on Bray-Curtis dissimilarity (Table S7). There was no evidence for effects of host sex or morphology on unweighted UniFrac (Table S8) or weighted UniFrac (Table S9) in any population. Next, we tested for correlations between beta diversity and morphological divergence among fish from the same lake. However, we found no evidence for such correlations in any of the populations and any of the three beta diversity metrics (Tables S10-12).

### Gut microbiota dissimilarity on the population level (across population beta diversity)

Next, we used multiple regression on distance matrices (MRM) to test the effects of divergence in free-living bacterial communities of the lakes, host morphology and genetics, and geographic distance on the three gut microbiota beta diversity metrics summarized on the population level. For Bray-Curtis dissimilarity, the model was significant (*F* = 12.379, *P* = 0.001) and explained 38.5% of the variation in gut microbiota beta diversity; host morphological divergence was the only significant predictor (*β* = 0.004, *P* = 0.001) (Figure 5A). For unweighted UniFrac, the model was not significant (*F* = 0.645, *P* = 0.947) and only explained 3.2% of the variation with no significant predictor variables. For weighted UniFrac, the model was significant (*F* = 11.437, *P* = 0.032) and explained 36.6% of the variation with divergence in free-living bacterial communities as the sole significant predictor (*β* = 0.169, *P* = 0.043) (Figure 5B).

**Figure 5:**
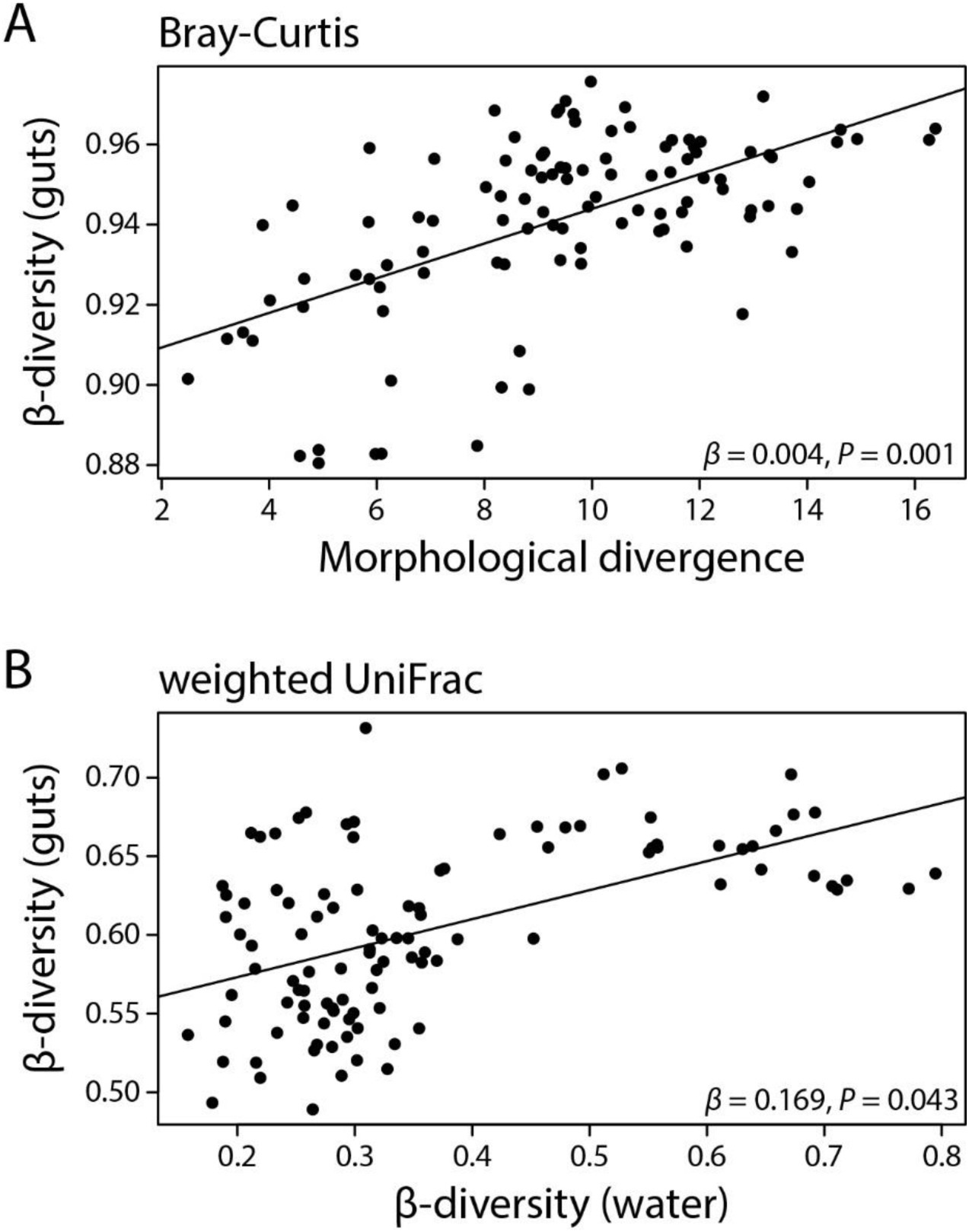
We used multiple regression on distance matrices (MRM) to find whether gut microbiota divergence (beta diversity) on the population is associated with divergence in free-living bacterial communities of the lakes, host morphology and genetics, as well as geographic distance. Population-level beta diversity was explained by host morphological divergence for Bray-Curtis dissimilarity (A) and by divergence in free-living bacterial communities for weighted UniFrac (B). No significant predictors were found for unweighted UniFrac.

## Discussion

This study focused on understanding the effects of host morphology (a proxy for trophic ecology), sex, genetic divergence, and environmental bacterial communities and physicochemical characteristics on gut microbiota composition and diversity across wild populations of threespine stickleback fish from 20 lakes on Vancouver Island, Canada. Across populations, we found substantial variation in the taxonomic composition and diversity of gut microbial communities (Figures 2 & 4). Further, alpha diversity was strongly affected by host sex (Figure 2C) and morphology (body shape), but not by environmental characteristics, suggesting a stronger effect of host traits compared to the conditions of the lake environment. Beta diversity was affected by both host traits and environmental characteristics but the factors included in our models explained only a very small proportion (<1%) of the variation in beta diversity. Multiple regression on distance matrices across populations further revealed that gut microbiota divergence among stickleback populations is associated with morphological divergence based on Bray-Curtis dissimilarity and with divergence of environmental bacterial communities based on weighted UniFrac metric. Yet, within-population analyses found limited evidence for associations between any host traits and gut microbiota alpha and beta diversity, with only a few populations showing significant effects. Overall, our study hints at the relative importance of host traits compared to environmental characteristics in shaping the stickleback gut microbiota across wild populations, with implications for understanding host-microbiota interactions in natural environments.

The core microbiota refers to a subset of microbial taxa that are consistently associated with a specific host, environment, or sample type. There are various approaches to defining and measuring the core microbiota (Shade and Handelsman 2012, Neu et al. 2021). Here, we defined ASVs that occurred in at least 50% or 80% of host individuals as part of the core microbiota. The results of our study indicate a weakly defined core gut microbiota, both within and across stickleback populations. The limited sharing of ASVs among hosts, with only 0.6% of hosts sharing any given ASV on average, highlights the high degree of individual specificity in gut microbial communities. At the population level, the number of shared ASVs varied greatly, with no population exhibiting a strongly defined core microbiota. For example, the number of ASVs shared by at least 50% of the population ranged from 4 to 16 across lakes, while no lake exhibited more than 3 ASVs shared among 80% of hosts. The overall weak evidence for a core microbiota is consistent with the high within-population beta diversity, indicating substantial variability in the gut microbiota even among fish within the same population (Figure 3B). This variation further underscores the complexity and individuality of microbial colonization within host populations, possibly driven by local environmental factors or host genetics. The presence of a small number of bacterial families (Halomonadaceae, Burkholderiaceae, Mycobacteriaceae) as part of the core microbiota across multiple populations suggests that these taxa may play important roles in the stickleback gut microbiota. Notably, the genus Burkholderia has been found to be part of the core microbiota of lake whitefish (Sevellec et al. 2018) and zebrafish (Roeselers et al. 2011), hinting at the importance of these bacteria for their fish hosts. Yet, while certain bacterial taxa may have a broad distribution, the stickleback gut microbiota is highly individualized, with host-associated and environmental factors potentially contributing to the observed variation. It is currently unclear how our results compare to other fish (or vertebrate) species, as measures and thresholds to define the core microbiota are inconsistent across studies (reviewed in Neu et al. 2021). The lack of a standardized framework makes it difficult to assess the degree to which different host species maintain a core microbiota and how taxonomically variable core microbiota are across species. A comprehensive meta-analysis would be valuable in addressing this issue, as has been done for reptiles (Hoffbeck et al. 2023), and could provide insights into the key factors shaping core microbiota across various hosts and environments.

One key observation from our study is the substantial influence of host population on gut microbiota diversity, suggesting that ecological variation across stickleback populations and adaptation to local conditions play pivotal roles in shaping the taxonomic composition and diversity of gut microbial communities. Previous work in stickleback found support for host diet (Bolnick et al. 2014b, Smith et al. 2015, Rennison et al. 2019, Härer and Rennison 2024) and genotype (Smith et al. 2015, Steury et al. 2019, Small et al. 2023) being important contributors to gut microbiota variation within and among populations. In our study, gut microbiota divergence among populations appeared to be driven by host morphological divergence (Figure 5A), but not by genetic divergence. In stickleback, body shape variation is associated with divergence in trophic ecology (Schluter 1993, Bell and Foster 1994) and, thus, our results suggest that the observed correlation between divergence in body shape and gut microbiota composition may be an effect of differences in host diet. However, in order to test this directly and draw more robust conclusions, diet information would need to be collected for individual hosts using stable isotope analysis or metabarcoding of gut contents (Nielsen et al. 2018). The lack of an association between genetic divergence and in gut microbiota composition was rather surprising but could be driven by overall low levels of genetic divergence among the populations included in our study, with F_ST_ values ranging from 0.026 to 0.091 among populations. In contrast, a study on stickleback from Oregon found an effect of genetic divergence on the gut microbiota but F_ST_ values ranged from 0.021 to 0.575 (Steury et al. 2019). Thus, we hypothesize that in our study the limited genetic divergence among the populations was insufficient to structure gut microbiota composition. The observed lack of association might also be due to the fact that genomic divergence data was obtained from a previous study (Bolnick and Ballare 2020) and, thus, come from different individuals than those for which we collected gut microbiota data. However, we believe this is improbable as genome-wide allele frequencies are unlikely to differ substantially from year to year, thus mean F_ST_ estimates should be little affected. It is worth emphasizing that despite this low genetic divergence, we observed high levels of beta diversity and gut microbiota structuring across populations. This may be further evidence of the role of trophic ecology driving patterns of gut microbiota differentiation, as prior work has shown that trophic divergence can arise in conjunction with low levels of genomic divergence (Bolnick and Ballare 2020, Härer et al. 2021) and we have evidence linking trophic morphology to gut microbiota diversity patterns. Despite limitations in our study, the results indicate that host body shape can be used to predict variation in stickleback gut microbiota diversity and composition. Hence, the observed association between these two factors underscores the potential eco-evolutionary implications of trophic specialization on host-associated microbial communities.

We detected mixed evidence for potential effects of specific host traits and environmental characteristics on gut microbiota alpha and beta diversity across stickleback populations. Host sex explained small but significant proportions of gut microbiota variation (beta diversity) and females had consistently higher alpha diversity compared to males across the three alpha diversity metrics used here (Figure 2C). In the literature, there is mixed evidence for sex-dependent differences in gut microbiota composition and diversity. Several studies in mammals found no or negligible effects of host sex on the gut microbiota of humans (Lay et al. 2005), ring-tailed lemurs (Bennett et al. 2016), mice (Kovacs et al. 2011), and North American red squirrels (Bobbie et al. 2017, Ren et al. 2017). In contrast, there are sex-specific differences of the gut microbiota in chimpanzees from Gombe National Park, which could be produced by differences in feeding behavior between females and males (Degnan et al. 2012). Previously, it has been found in stickleback that diet had sex-dependent effects on the gut microbiota within one population (Bolnick et al. 2014c). However, no effects of host sex on the gut microbiota were found across two other populations and their hybrids (Small et al. 2023), across ten populations from lakes, rivers and estuaries on Vancouver Island, Canada (Smith et al. 2015) or in benthic and limnetic ecotypes reared in experimental ponds (Härer et al. 2024). Thus, our study is the first to provide compelling evidence for sex-dependent differences in gut microbiota alpha and beta diversity across stickleback populations. Yet, it currently remains unknown whether the observed gut microbiota differences are produced by sexual dimorphism in trophic ecology or in physiology (immune system function, hormone levels, etc.), which opens future lines of research to investigate the mechanisms that produce these sex differences in stickleback fish.

Notably, there was an association between divergence in gut microbial and free-living bacterial communities based on the weighted UniFrac metric (Figure 5B), suggesting that the stickleback gut microbiota is to some extent affected by the bacterial communities of the lake environment. Yet, we found distinct clustering according to sample type (fish guts vs lake water; Figure 4), which is in line with results from previous studies showing that gut microbial communities of fish are often highly distinct from their free-living counterparts of the aquatic environment (Sevellec et al. 2018, Härer et al. 2020, Sadeghi et al. 2023). Microbes from the environment, particularly through ingested water and food items, are frequently introduced into the stickleback gut, where they may either pass through without establishing, fail to survive the local conditions in the gut, or successfully colonize and become part of the resident microbiota. Indeed, a previous study in stickleback showed that a substantial proportion of bacterial lineages found in the gut microbiota may be acquired from the aquatic environment (Smith et al. 2015). Thus, environmental bacteria appear to contribute to the assembly of the stickleback gut microbiota but it remains to be determined whether the association between gut-associated and free-living bacterial communities is produced by fish picking up and incorporating environmental bacteria into their gut microbiota or by shared environmental conditions. We argue it is unlikely that the observed correlation between gut-associated and free-living bacterial communities is driven by geographically structured environmental variation because the geographic distance between lakes did not affect the stickleback gut microbiota. If environmental gradients were to shape both free-living and host-associated bacterial communities, a higher similarity of environmental conditions (e.g., temperature and precipitation) might be expected between lakes in close proximity. A decrease in compositional similarity among bacterial communities with geographic distance (i.e., distance-decay of similarity) (Nekola and White 1999) has been found in free-living bacterial communities from various habitats and geographic regions (Martiny et al. 2006, Martiny et al. 2011, Clark et al. 2021). Yet, the apparent lack of an effect of geographic distance in our study system suggests that distance-decay of similarity might not necessarily apply to host-associated gut microbial communities. In sum, while the stickleback gut microbiota is distinct from free-living bacterial communities, environmental bacteria appear to play a significant role in shaping its composition, highlighting the complex interactions between host, gut, and environment.

While several host traits and environmental characteristics had small but significant effects on gut microbial communities across host populations (beta diversity), our within-population analyses revealed much weaker patterns. Across all alpha and beta diversity metrics, we observed significant associations between either host sex or morphology and the gut microbiota for a maximum of two lakes per metric, which implies non-consistency of such associations across stickleback populations. Regarding host sex, it remains to be answered why some populations show gut microbiota differences between males and females while others do not, but population-specific differences in sexual dimorphism of diet and trophic traits could be involved (McGee and Wainwright 2013, Bolnick et al. 2014c, Reimchen et al. 2016). The lack of associations between host morphology and the gut microbiota in most populations could be explained by low levels of divergence among host individuals of the same population compared to the extent of divergence among individuals of different populations. This interpretation is supported by the observation that host population had a much larger effect on gut microbiota composition than host sex or morphology. Thus, our results suggest that in our study system of 20 stickleback lake populations from Vancouver Island, Canada, the gut microbiota is mainly structured across, rather than within, populations.

## Conclusions

Our study underscores the multifaceted nature of host-microbiota interactions and the importance of considering both host-associated and environmental factors in unraveling gut microbiota diversity across wild host populations. Notably, we found strong divergence in gut microbiota composition among closely related populations with minimal genetic differentiation but substantial variation in trophic ecology. This finding suggests that dietary differences may drive substantial microbiota differentiation across populations, though the role of other environmental factors, such as local abiotic conditions and biotic interactions, need further attention. While our study provides valuable insights, future research should aim to integrate morphological, dietary, and genetic analyses alongside gut microbiota characterizations within the same host individuals to deepen our understanding of mechanistic underpinning of these interactions. Despite these limitations, the insights gained from our research contribute to the growing body of knowledge on microbial ecology and can have implications for our understanding of the eco-evolutionary dynamics of host-associated microbial communities. By providing evidence for the role of trophic ecology in shaping microbiota divergence, this study reinforces the importance of exploring eco-evolutionary dynamics of host-microbiome interactions across diverse ecological contexts.

## Declarations

### Ethics approval and consent to participate

Samples were collected in 2020, 2021, and 2022 under British Columbia Fish Collection permits NA20-602264, MRVI21-619908, and NA22-713085, respectively.

### Authors’ contributions

A.H., E.K., and D.J.R conceptualized the study. E.K. prepared and measured fish specimens, A.H. conducted the gut microbiota library preparation and analyzed the data. A.H. wrote the manuscript with input from E.K. and D.J.R.

## Acknowledgements

We thank Tristan Kosciuch, Carolyn McKinnon, and Christian Carson for assistance with collecting samples from the field and Camila Arregui, Christine Frazier, Alina Shahin, Ashley Toba, Mia Tonkin, and Isabella Villalobos-Arriaga for collecting morphological data.

